# Intracellular trafficking of furin enhances cellular intoxication by recombinant immunotoxins based on *Pseudomonas* exotoxin A

**DOI:** 10.1101/2023.05.31.542721

**Authors:** Brian D. Grossman, Jack D. Sanford, Yuyi Zhu, Cynthia B. Zeller, John E. Weldon

## Abstract

Furin is a mammalian serine protease with important roles in cellular homeostasis and disease. It cleaves and activates numerous endogenous and exogenous substrates, including the SARS-CoV-2 viral spike protein and protein toxins such as diphtheria toxin and *Pseudomonas* exotoxin A (PE). Recombinant immunotoxins (RITs) are toxin conjugates used as cancer therapeutics that connect tumor-directed antibodies with toxins for targeted cell killing. RITs based on PE have shown success in treating a variety of cancers, but often suffer from safety and efficacy concerns when used clinically. We have explored furin as a potential limiting factor in the intoxication pathway of PE-based RITs. Although the furin has widely recognized importance in RIT intoxication, its role is incompletely understood. Circumstantial evidence suggests that furin may act as a transporter for RITs in addition to its role of activation by cleavage. Here, we describe the creation of a CRISPR-engineered furin-deficient HEK293 cell line, ΔFur293. Using ΔFur293 and derivatives that express mutant forms of furin, we confirm the importance of furin in the PE RIT intoxication pathway and show that furin trafficking has a significant impact on RIT efficacy. Our data support the hypothesis that furin acts as a transporter during RIT intoxication, and suggest furin as a target to improve the effectiveness of RITs.

## Introduction

Furin (EC 3.4.21.75) is a ubiquitously expressed mammalian protease that belongs to the *proprotein convertase subtilisin/kexin* (PCSK) family of serine endoproteases (reviewed by Seidah and Prat^1^). Within the secretory pathway, furin proteolytically activates numerous proprotein substrates that include endogenous pro-hormones and growth factors as well as pathogen-associated viral glycoproteins and bacterial toxins. Furin is important for maintaining cellular homeostasis and has an essential role in embryogenesis, It is also implicated in diseases ranging from Alzheimer’s and cancer to bird flu, Ebola, and anthrax (reviewed by Thomas^2^).

Recently, furin has been implicated in viral infection by SARS-CoV-2, and there is evidence that a novel furin cleavage site on the viral spike protein may be responsible for its high infectivity and overall virulence^3–5^.

Studies describing the role of furin in both normal cellular activity and disease have provided important molecular insights (reviewed in several publications^2,6,7^). Evaluation of furin knockout mice showed an embryonic lethal phenotype^8^, but furin knockdown by siRNA was not lethal to cultured cells^9^. Furin expression varies widely among the cell lines evaluated in the Human Protein Atlas (www.proteinatlas.org)^10^, but naturally furin-deficient cell lines, such as the CHO-derived RPE.40^11,12^ and colon carcinoma line LoVo^13,14^, are rare. The limited selection of furin-deficient cell lines and variation among existing lines has limited our capability to study the role of furin in cultured cells.

One exogenous furin substrate is *Pseudomonas* exotoxin A (PE), a bacterial toxin secreted by *Pseudomonas aeruginosa*. PE can be conjugated to antibody fragments to make a targeted cell killing agent called a recombinant immunotoxin (RIT) (reviewed by Weldon and Pastan^15^). RITs are used therapeutically for the treatment of cancers; several are in clinical trials and one (moxetumomab pasudotox^16^) has been FDA-approved for the treatment of hairy cell leukemia. Furin plays an important, but incompletely understood, role during the intoxication of PE and PE-based RITs. Prior research has established that furin proteolytically activates the toxin during its intracellular trafficking route^17–20^. In addition, furin-deficient LoVo cells show poor sensitivity to RITs^21^, furin inhibitors diminish RIT activity^22^, and mutations to the furin cleavage site of RITs can influence their cellular toxicity^22^.

In addition to furin-mediated cleavage, circumstantial evidence suggests that furin may act as a transporter for PE and PE-based RITs. The furin cleavage site on PE is a remarkably poor substrate for furin, demonstrating the most unfavorable cleavage characteristics of all substrate sequences evaluated in two separate studies^23,24^. Our own unpublished experience with *in vitro* cleavage assays confirms this. Mutations around the furin cleavage site of RITs can enhance both furin cleavage and RIT cytotoxicity, but there was no correlation between the two parameters^22^. Evidence also suggests that furin can act as a non-proteolytic chaperone, as demonstrated in the case of matrix metalloproteinase-28 (MMP-28) transport from the TGN to the cell surface, where MMP-28 is released without being cleaved^25^. Taken together, these points suggest that the role of furin is more complex than a cleavage activation step in the intoxication pathway.

In this study, our objective was to investigate the role of furin in the intoxication pathway of PE-based RITs. Specifically, we sought to understand the relationship between the cleavage, cytotoxicity, and intracellular localization of RITs and furin. We hypothesized that in addition to cleavage, furin can also function in directing RITs to the necessary subcellular compartments required for productive intoxication, such as from endosomes to the Golgi..

We first describe the generation and characterization of a furin-deficient HEK293 cell line, ΔFur293. HEK293 cells are well suited for understanding the role of furin in cellular processes; they have excellent growth characteristics, are widely studied, and express clearly detectable levels of furin. We employed a variant HEK293 line, HEK293 FRT, that contains a single Flp recombinase recognition target site (FRT) stably integrated into its genome. Using a CRISPR/Cas9 system, we genetically modified HEK293 FRT cells to eliminate endogenous furin expression. Results were confirmed by sequencing, western blots, cytotoxicity assays, intracellular cleavage assays, and complementation with transgenic furin. The ΔFur293 line will be a convenient and valuable tool for studies evaluating the role of furin in a variety of natural and disease-related processes.

We then employed the ΔFur293 cell line to investigate the influence of furin on the activity of PE-based RITs. We stably introduced mutant forms of furin with either altered intracellular trafficking patterns or decreased catalytic activity into ΔFur293 cells. The results of RIT cytotoxicity assays support the importance of furin in the intoxication pathway of PE-based RITs, and indicate that both furin cleavage and trafficking play an important role during RIT intoxication.

### Experimental procedures Design of sgRNAs

Single guide RNAs (sgRNAs) to the furin gene were designed using the Broad Institute sgRNA design tool (CRISPick)^26,27^. Three sgRNAs that scored highly and targeted locations early in the furin gene were selected: GCGCATCCCTGGAGGCCCAG (score 0.735013), GAAGGTCTTCACCAACACGT (score 0.606152), and GCTACCACCCATAGCAACCA (score 0.689511).

#### Primers and plasmid constructs

All oligonucleotides used in this study obtained from Integrated DNA Technologies, Inc. (IDT, Coralville, IA) and are detailed in Supplementary Table 2. Oligonucleotides corresponding to the selected furin sgRNAs were ligated into the pSpCas9(BB)-2A-Puro (PX459) V2.0 plasmid, a gift from Dr. Feng Zhang (Addgene plasmid #62988; http://n2t.net/addgene:62988; RRID:Addgene_62988)^28^. Human furin cDNA (product #RC204279) was obtained from OriGene (Rockville, MD) and inserted into the pcDNA5/FRT plasmid for stable transfection into HEK293 FRT cells. Mutations in the furin gene was performed by site-directed mutagenesis.

The HB21-LR expression plasmid pHB21-LR was generated by mutation of the original pHB21 plasmid from the Pastan Lab. Plasmids were transformed into NEB5α competent *E. coli* (NEB, Ipswich, MA) and DNA was harvested using the Nucleospin plasmid isolation kit (Macherey Nagel, Bethlehem, PA). All constructs were confirmed by gel electrophoresis and sequencing (Macrogen USA, Rockville, MD).

All restriction enzymes were purchased from NEB and PCR amplification was performed with Q5 High-Fidelity PCR (NEB) following the manufacturer’s recommendations. DNA ligations were performed using the Blunt/TA Ligase Master Mix (NEB) followed by transformation into NEB5α competent *E. coli* cells.

#### Mammalian cell culture

HEK293 FRT cells were a kind gift from Dr. Ira Pastan (NCI/NIH). They contain a single stably integrated Flp recombinase recognition target (FRT) site at a transcriptionally active genomic locus. LoVo cells (CCL-229) were obtained from ATCC (Manassas, VA). All cells were grown at 37ºC with 5% CO_2_ in high glucose Dulbecco’s Modified Eagle Medium (DMEM) with sodium pyruvate (Corning) supplemented with 10% FBS (GeneMate) and 2 mM Glutagro

L-glutamine supplement (Corning). HEK293 FRT cells were cultured in zeocin (100 μg/ml, Invitrogen) to maintain the integrated FRT site. Cells were passaged once they reached >80% confluency.

Transfections were performed using Turbofection 8.0 (Origene, Rockville, MD) following the manufacturer’s recommendations. Individual sgRNA plasmids and combinations of plasmids were transfected. Puromycin was added to each well at a concentration of 3 μg/ml following initial passaging. The transfection utilizing all three sgRNA plasmids showed the most marked reduction in furin levels as determined by western blot analysis and was chosen for clonal selection by serial dilution in a 96-well plate according to protocol^29^.

#### Total cell lysates

Cells at >80% confluency were harvested and resuspended at a density of 5x10^6^ in 1 ml of cold RIPA buffer (50 mM Tris-HCl pH 8, 150 mM NaCl, 1 mM EDTA, 1% Triton X-100, 0.5% sodium deoxycholate, 0.1% SDS) with a broad-spectrum protease inhibitor (Thermo Scientific: A32953). The mixture was incubated on ice for 10 minutes, sonicated, and stored at - 20°C. Protein concentrations were determined using a bicinchoninic acid assay (BCA) protein assay kit (Thermo Scientific: 23227).

#### Genomic DNA sequencing

Genomic DNA was extracted from the selected monoclonal population (Wizard® Genomic DNA Purification Kit, Promega, Madison, WI) following the manufacturer’s recommended protocol. DNA was PCR amplified (primers hFUR Exon 1-3 PCR For and hFUR Exon 1-3 PCR Rev) and purified (Monarch PCR & DNA cleanup kit, NEB). The resulting PCR product was sequenced using the hFur Exon 1 Seq primer.

To confirm the genomic sequence of ΔFur293, overlap extension PCR was performed on the genomic DNA to clone exon 2 of the furin gene into the pUC57^30,31^. Briefly, genomic DNA from ΔFur293 cells was PCR amplified (primers hFUR Exon 1-3 PCR For and hFUR Exon 1-3 PCR Rev). This PCR product was purified, and a 2 ng/μL solution was prepared. A second PCR reaction was performed using the product from the reaction 1 and primers that contain overhangs complementary to portions of pUC57-kan (pUC57 Furin Ex 2 OvEx C+A and pUC57 Furin Ex 2 OvEx D+B). The resulting product from reaction 2 was purified and used as a primer for PCR amplification of the pUC57 plasmid. The product from reaction 3 was *Dpn*I treated, purified, and transformed into NEB5α. Multiple colonies were grown for DNA extraction. In total, 10 plasmids were sequenced using the M13 R primer.

### Recombinant immunotoxin purification

The recombinant immunotoxin HB21-LR^32^, derived from the single-chain anti-transferrin receptor RIT HB21^33^ using the “LR” modifications^34^, was expressed and purified following the published protocol^35^, except for the following modifications. A starting quantity of 40 mg was used instead of 100 mg. All reagents were scaled proportionally, but concentrations remained equivalent. The column chromatography employed a single anion-exchange step instead of the two-step protocol. In place of the Q Sepharose and Mono Q columns as described^35^, only a HiTrap Q HP column (Cytiva, Marlborough, MA) was used. The expression plasmid was a kind gift from Dr. Ira Pastan.

### Recombinant immunotoxin internalization and cleavage assay

Cells (10^6^) were seeded in 60x15 mm dishes with 4 mL fresh medium and incubated for approximately 24 hours. After reaching 60-80% confluency, HB21-LR was added directly to the 4 mL of media to a final concentration of 0.1 μg/mL. Cells were incubated at 37°C for various time intervals, after which the media was aspirated and the dish was washed consecutively with 1 mL cold DPBS, 1 mL cold stripping buffer (1 mg/mL BSA in 0.2 M glycine, pH 2.5), and 1 mL cold DPBS. Ice-cold RIPA buffer (100 μL) with protease inhibitor cocktail (ThermoFisher) was added to the dish, cells were dislodged using a cell scraper, and the suspended cells were transferred to a microcentrifuge tube. This process was repeated with an additional 100 μL of ice-cold RIPA buffer to collect any cells remaining on the dish. The RIPA solution was then used to prepare total cell lysates as previously described.

#### Western Blots

Cell lysates containing 20 μg of total protein were analyzed by western blot. The specific primary and secondary antibodies and their corresponding dilutions are indicated in the appropriate figure legends. For furin expression level analysis, mouse anti-furin antibody (mAb, Santa Cruz Biotechnology: sc-133141), mouse anti-β-actin (mAb, ThermoFisher Scientific: BA3R), and goat anti-mouse IgG alkaline phosphatase conjugated antibody (mAb, Santa Cruz Biotechnology: sc-2058) were used. For internalization and cleavage assay analysis, rabbit anti-PE (pAb, NCI/NIH), rabbit anti-β-actin (mAb, Boster Bio: M01263), and mouse anti-rabbit IgG alkaline phosphatase conjugated antibody (mAb, Santa Cruz Biotechnology: sc-2358) were used. Antibodies were visualized chromogenically by incubating the membrane in an alkaline phosphatase buffer (130mM Tris, pH 9, 150 mM NaCl, 100 μM MgCl_2_) and 2% BCIP/NBT (Roche: 11681451001).

Densitometry was performed using Bio-Rad’s Image Lab Software (Version 6.01). For expression level analysis, the ratio of the target intensity to that of the loading control was calculated for each sample and normalized to the cell line with the lowest expression. For internalization and cleavage analysis, the ratio of the intensity between total internalized toxin and cleaved toxin was calculated.

#### Cytotoxicity assays

Viability of cell lines treated with immunotoxins was measured using the Cell Counting Kit-8 assay (Dojindo Molecular Technologies: CK04) as previously described^22^. PPCI was purchased from Sigma-Aldrich (#537076). Comparisons between two groups were analyzed using a paired, two-tailed t-test assuming a Gaussian distribution. Comparisons of three or more groups were analyzed using a one-way ANOVA with a Geisser-Greenhouse correction and a Tukey-Kramer multiple comparisons test using GraphPad Prism.

#### STR profiling

Genomic DNA was extracted from cultured cells using the Wizard® Genomic DNA Purification Kit (Promega) and amplified using the Investigator 24plex QS Kit (Qiagen, Hilden, Germany) on a Veriti 96-well thermal cycler (Applied Biosystems, Foster City, CA). Each sample was separated on a 3500 Genetic Analyzer (Applied Biosystems). The resulting electropherograms were analyzed with GeneMapper ID-X software (Applied Biosystems) version 1.5.

## Results

### Furin knockout

Using CRISPR-Cas9 based gene editing, we generated a furin deficient HEK293 FRT cell line, ΔFur293. In brief, three single guide RNAs (sgRNAs) were designed that targeted exon 2 of the human furin gene. HEK293 FRT cells were transfected with genes encoding Cas9 and all three sgRNAs concurrently. Surviving cell populations were evaluated for furin expression by western blot (Figure 1A), and one population (#2) was subcloned by serial dilution. The subcloned populations were evaluated for furin expression by western blot (Figure 1B). One clone (#1) without visible furin expression was selected and designated as ΔFur293 for further analysis.

**Figure 1.**
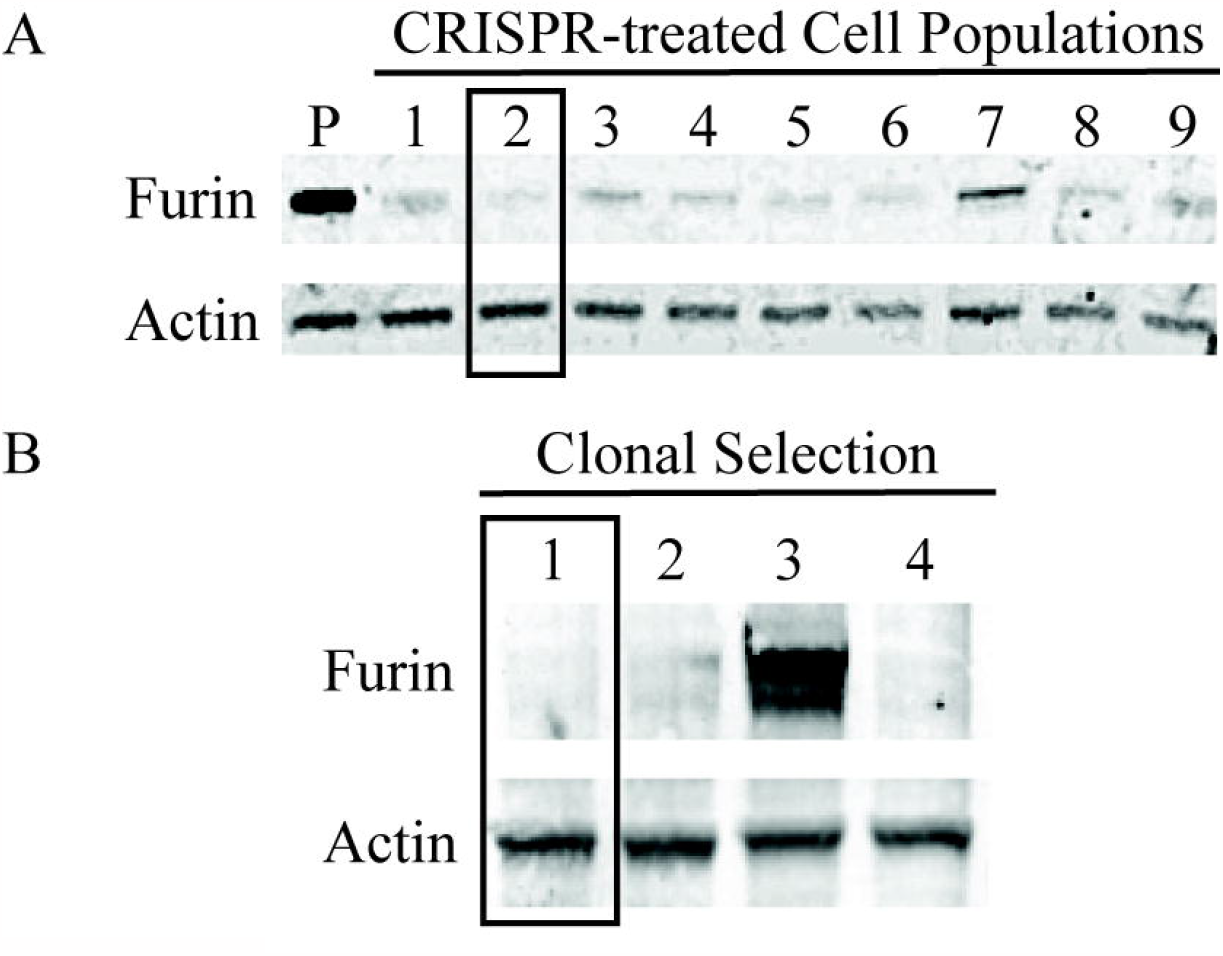
Elimination of furin expression from HEK293 cells. HEK293 FRT cells were transfected with plasmids containing the genes coding for Cas9 and three furin sgRNAs. Lysates from nine surviving cell populations were evaluated for furin expression by western blot (panel A). The untreated parental (P) HEK293 FRT line is also shown for comparison. Cells from well 2 (boxed) were clonally selected by serial dilution in a 96-well plate. Lysates from four clones were evaluated for furin expression by western blot (panel B). Well 1 (boxed) was selected for further analysis.

To confirm the mutation in the furin gene, a 1300-bp section of the gene was amplified using genomic DNA extracted from ΔFur293 cells and directly sequenced. Initial results showed that exon 2 of the furin gene had a deletion of approximately 105 nucleotides. However, the sequencing chromatogram indicated heterogeneity among the alleles of the furin gene and prevented an exact sequence. To precisely determine the sequence of each allele, the gene segment was cloned into a pUC57-kan plasmid using overlap extension PCR, allowing for precise sequencing of each clone. Sequencing of 10 clones resulted in near equal proportions of two distinct alleles differing by a single nucleotide.

Aligning the sequences to the human furin gene (Ensemble Transcript ID: ENST00000268171.8) showed one allele with a 105 bp deletion from nucleotides 7168-7272 and a second allele with a 106 bp deletion from nucleotides 7168-7273 (Supplementary Figure 1). The first allele (105 bp deletion) results in the loss of 34 amino acids at positions 7-41 in the furin protein sequence. The deletion spans most of the signal peptide domain (residues 1-26) and the amino terminal portion of the prodomain (residues 27-107). The second allele (106 bp deletion) results in a similar loss of amino acids 7-41 and also introduces a frameshift mutation beginning at position 42, leading to a nonsense product.

#### STR profile

To confirm the identity and establish a baseline STR profile for our cell lines, we evaluated 22 loci from ΔFur293 and its parent cell line HEK293 FRT. We compared our results to each other and to a reference profile for HEK293 (RRID:CVCL_0045) in the *Cellosaurus* database^36^. These results are shown in Supplementary Table 1. HEK293 FRT and ΔFur293 were identical to each other at 20 of the 22 loci and showed a partial match at 2 loci (D19S433 and D8S1179).

Six of the loci we evaluated were not reported in the *Cellosaurus* database and a comparison to the reference profile could not be made. Variations in the reference profile were reported at four loci in the *Cellosaurus* database, and a search of published literature and biological resource centers found variations at an additional four loci. Both HEK293 FRT and ΔFur293 were consistent with one of the reference variations at these eight loci. Seven of the remaining eight loci were identical among all three profiles. One locus (D8S1179) showed a partial match with HEK293 FRT.

#### Cytotoxicity Analyses

To evaluate the effect of furin on the efficacy of recombinant immunotoxins (RITs), the cytotoxicity of HB21-LR (anti-transferrin receptor single-chain variable fragment PE24 RIT)^32^ was evaluated against HEK293 FRT and ΔFur293 cells (Figure 2). Each cell line was evaluated in the presence and absence of proprotein convertase inhibitor (PPCI), a cell permeable reversible inhibitor of furin (K_i_=16 pM) that has been shown to have minimal toxicity at concentrations of up to 50 μM^37^. Preliminary analysis of PPCI indicated that a 1 μM concentration was adequate to achieve the maximum influence on RIT cytotoxicity (Supplementary Figure 2). A representative set of cytotoxicity assays is shown in Supplementary Figure 3.

**Figure 2.**
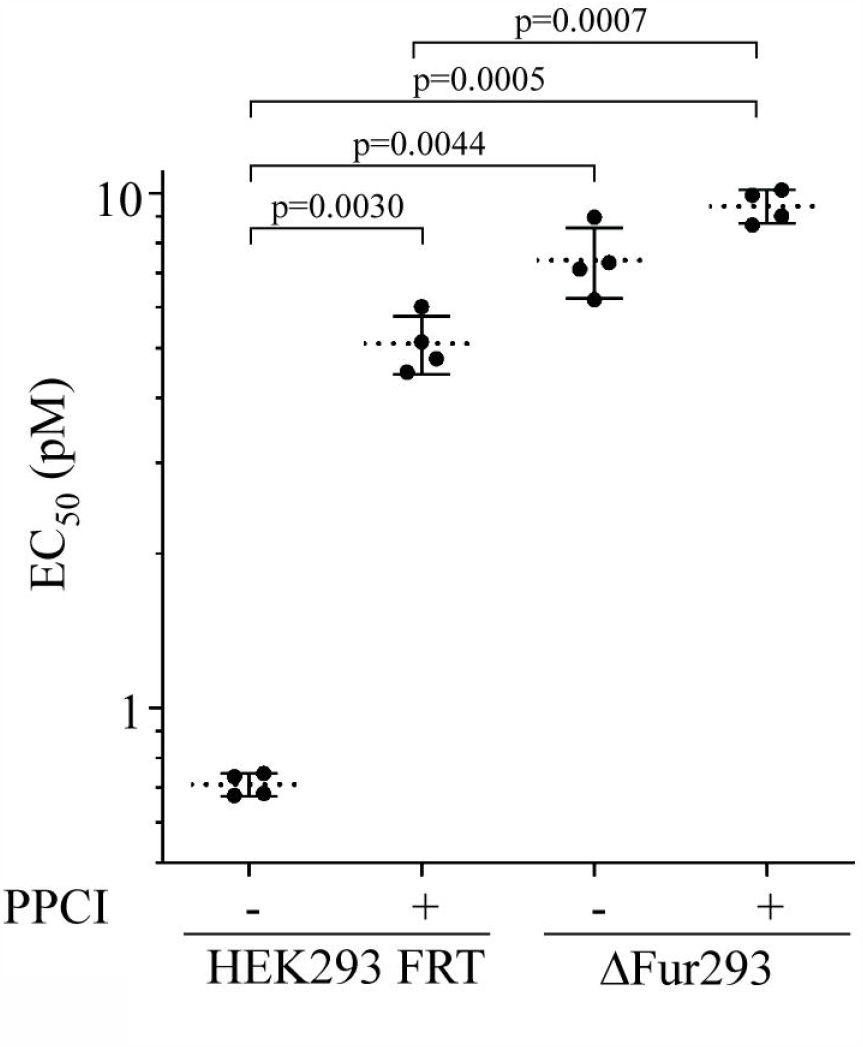
Cytotoxicity assays. HEK293 FRT and ΔFur293 cells were treated with the anti-transferrin receptor/PE24 RIT HB21-LR in the presence and absence of 1 μM PPCI furin inhibitor. EC_50_ (pM) values from four separate paired assays for each condition are plotted. Dotted lines denote the average value for each condition and error bars indicate the standard deviation. Significant p values (p<0.05) are reported from a one-way ANOVA performed as described. A representative cytotoxicity assay is shown in Supplementary Figure 3.

Analysis showed that the EC_50_ of HB21-LR against HEK293 FRT cells significantly increased by approximately 8-fold (p=0.0030) in the presence of PPCI. No difference in HB21-LR cytotoxicity against ΔFur293 cells with and without PPCI was found. When compared to untreated HEK293 FRT cells, ΔFur293 cells in the presence and absence of PPCI showed significant increases in EC_50_ of approximately 13-fold (p=0.0005) and 10-fold (p=0.0044).

Comparison of PPCI-treated HEK293 cells with ΔFur293 cells showed no statistically significant difference in EC_50_. Interestingly, a small (1.5-fold) but significant difference was observed between the EC_50_ of HEK293 FRT and ΔFur293 cells when both lines were in the presence of PPCI (p=0.0007). Overall, these data show that furin deletion results in dramatic decrease in HB21-LR toxicity.

As an additional control, we evaluated HB21-LR against the human colon carcinoma cell line LoVo, which lacks active furin^13,14^. As with ΔFur293 cells, no significant difference was observed with or without PPCI (Supplementary Figure 4).

**Figure 3.**
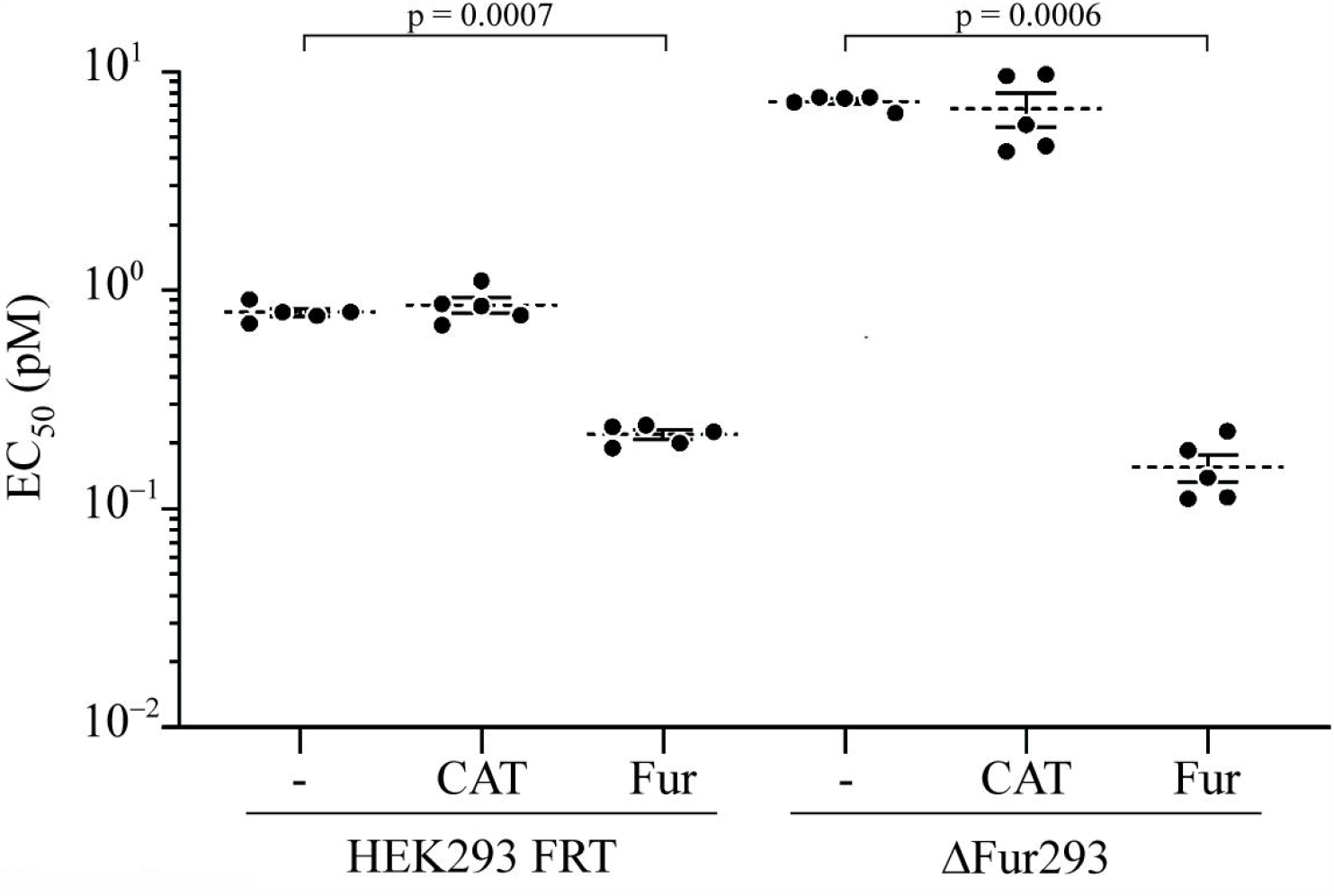
Furin complementation. HEK293 FRT and ΔFur293 cells were stably transfected with the gene for furin (Fur) or chloramphenicol acetyltransferase (CAT). All cell lines were then treated with the anti-transferrin receptor/PE24 RIT HB21-LR and evaluated for cytotoxicity. EC_50_ (pM) values from five separate paired assays for each line are plotted. Dotted lines denote the average value for each condition and error bars indicate the standard error. Significant p values (p<0.05) are reported from a one-way ANOVA performed as described.

**Figure 4.**
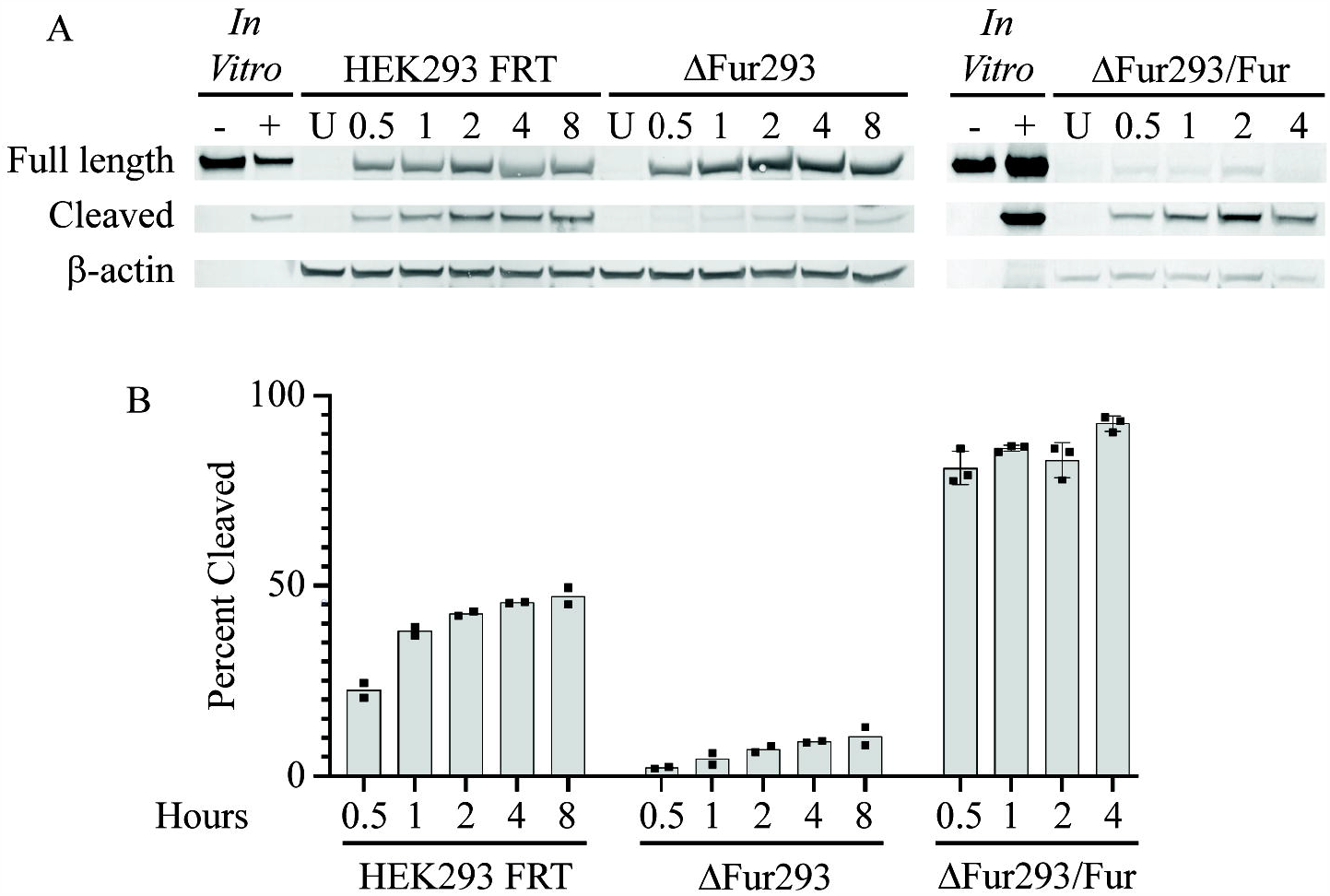
Cleavage assays. HEK293 FRT cells, ΔFur293 cells, and ΔFur293 cells stably expressing transgenic wild-type furin (ΔFur293/Fur) were incubated for various time intervals from 0.5 to 8 hours in culture with the anti-transferrin receptor/PE24 RIT HB21-LR. Whole cell lysates were evaluated for full length and cleaved HB21-LR by western blot (panel A) and densitometry (panel B) as described. Also shown are untreated (U) cell lysates for each cell line, HB21-LR with (+) and without (-) furin treatment in vitro, and the β-actin loading control. The ratio between the furin-cleaved band intensity and the total intensity of all RIT bands at each time point is plotted in panel B. The individual densitometric analysis values (points) and mean (bar) for at least two separate assays of each cell line are shown.

### Furin complementation

We next assessed the effect of restoring furin to ΔFur293 cells by stably transfecting them with the wild-type furin gene (Fur) inserted into the genomic FRT site (ΔFur293/Fur cells). Results from cytotoxicity assays are shown in Figure 3. As a control, we also evaluated ΔFur293 cells stably transfected with the gene for chloramphenicol acetyltransferase (CAT) at the FRT site (ΔFur293/CAT).

We then evaluated the cleavage efficiency of HB21-LR in each cell line over an 8 hour time course (Figure 4). Lysates from treated cells were examined for full length and cleaved HB21-LR by western blot at various time intervals (Figure 4A). Each band was evaluated by densitometry and normalized against the actin loading control. The cleaved fraction of total protein was plotted against time for each of the cell lines (Figure 4B). In HEK293 FRT cells, the percentage of total toxin cleaved within the cell peaked at 50%, while in ΔFur293 cells the percentage of cleaved toxin peaked at less than 10%. The ΔFur293/Fur cells demonstrated enhanced cleavage of the toxin, reaching a peak level of over 90% RIT cleaved at the 4 hour time point. We were unable to harvest a sufficient quantity of intact ΔFur293/Fur cells to analyze at the 8 hour time point due to their high sensitivity to HB21-LR.

#### Furin mutants

We next generated three mutants of furin designed to evaluate its role in RIT intoxication: two mutants with altered intracellular trafficking patterns (S773A/S775A and S773D/S775D) and one mutant with diminished catalytic activity (N295A). The trafficking mutants target two serine residues (positions 773 and 775) in the acidic cluster/TGN sorting signal within the cytoplasmic tail of furin. The phosphorylation state of these serine residues controls the trafficking of furin between the TGN and the cell surface^38^. The catalytic activity of furin cannot be completely eliminated from the mature protein because of its requirement for autocatalytic processing^39^. Reduced activity, however, can be obtained by mutation of an asparagine at position 295, which corresponds to the position of the oxyanion hole in its catalytic mechanism^40^.

The three mutants were stably transfected into the FRT site of ΔFur293 cells (ΔFur293/Fur^ADA^, ΔFur293/Fur^DDD^, ΔFur293/Fur^Ala-295^). Initial cytotoxicity analysis of these mutants showed significant differences between clones expressing the same furin variant (data not shown), which suggested that clones might exhibit variable expression levels of the furin transgene. To test this, we compared furin expression from two clones of each mutant, as well two clones of wild-type furin stably transfected into both ΔFur293 and HEK293 FRT cells.

Supplementary Figure 5B shows a western blot of cell lysates from each line. The results were quantified by densitometry (Supplementary Figure 5A) and used to normalize the results of cytotoxicity assays. This normalization eliminated discrepancies between different clones expressing the same mutant (Figure 5).

**Figure 5.**
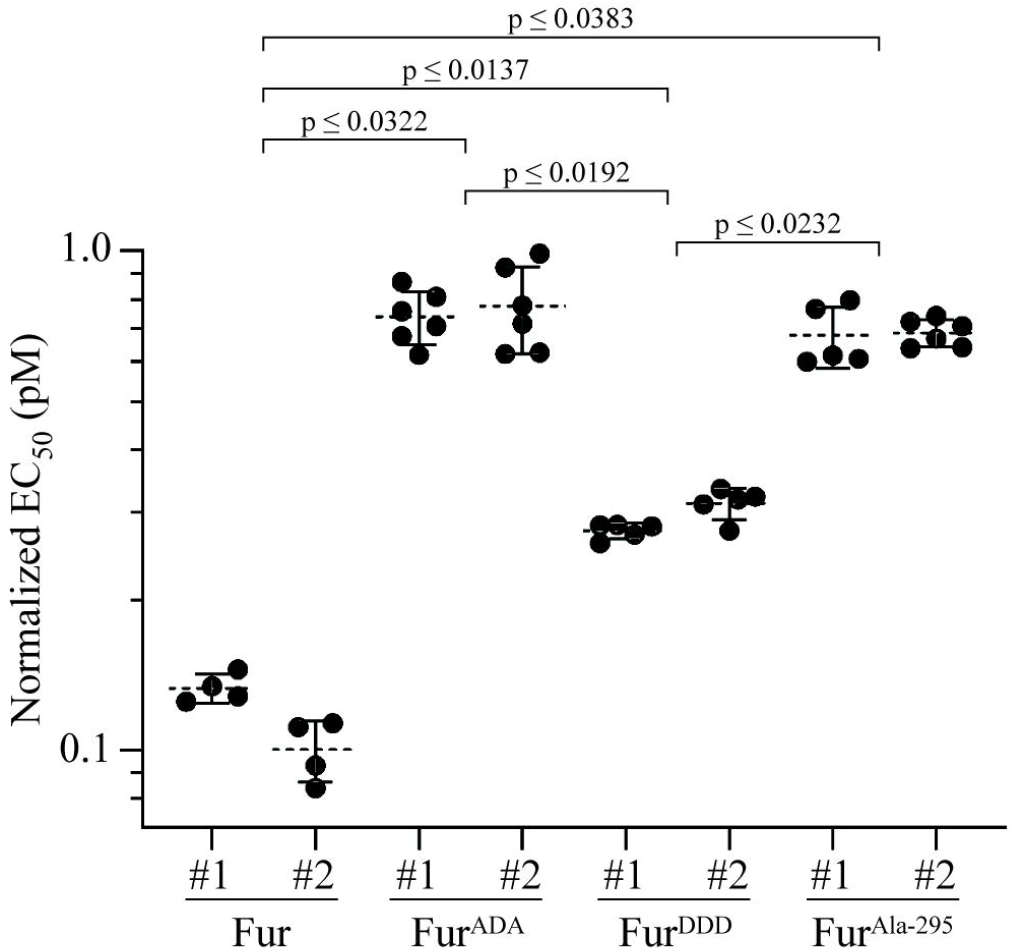
Complementation with mutant furin. ΔFur293 cells were stably transfected with genes for furin that contained mutations designed to impair its intracellular trafficking or catalytic function. Mutations S773A/S775A (ADA) and S773D/S775D (DDD) alter furin trafficking, while the N295A mutant (Ala-295) inhibits catalytic activity. Two separate clonal lineages stably transfected with mutant or wild-type furin were treated with the anti-transferrin receptor/PE RIT HB21-LR to assess cytotoxicity. The EC50 (pM) values from at least four separate assays for each line were normalized for furin expression levels and plotted. Dashed lines denote the average value for each clone and error bars indicate the standard error. The largest significant p values (p<0.05) between sets of clones from a one-way ANOVA performed as described are indicated. All p values are reported in Supplementary Table 3.

The S773A/S775A alanine trafficking mutant of furin (Fur^ADA^) causes it to become trapped in early endosomes^38,41^. We observed no difference between HB21-LR cleavage in ΔFur293/Fur and ΔFur293/Fur^ADA^ cells (Figures 4 and 6), and an 8-fold resistance to HB21-LR cytotoxicity (Figure 5). The S773D/S775D aspartic acid trafficking mutant of furin (Fur^DDD^) is depleted from early endosomes and preferentially retrieved to the TGN or cell surface, exhibiting a more disperse localization in the cell^38^. Cleavage was decreased by approximately 20% when compared to ΔFur293/Fur cells (Figures 4 and 6). Cytotoxicity was decreased nearly 3-fold compared to ΔFur293/Fur cells and increased 2.6-fold compared to ΔFur293/Fur^ADA^ cells (Figure 5). The N295A furin oxyanion hole mutant (Fur^Ala-295^) has decreased catalytic activity compared to wild type furin^40^ while having an identical trafficking pattern^42^. The ΔFur293/Fur^Ala-295^ cells are approximately 40% less efficient at cleaving HB21-LR when compared to ΔFur293/Fur cells (Figures 4 and 6). This resulted in an approximately 8-fold decrease in cytotoxicity relative to ΔFur293/Fur (Figure 5).

**Figure 6.**
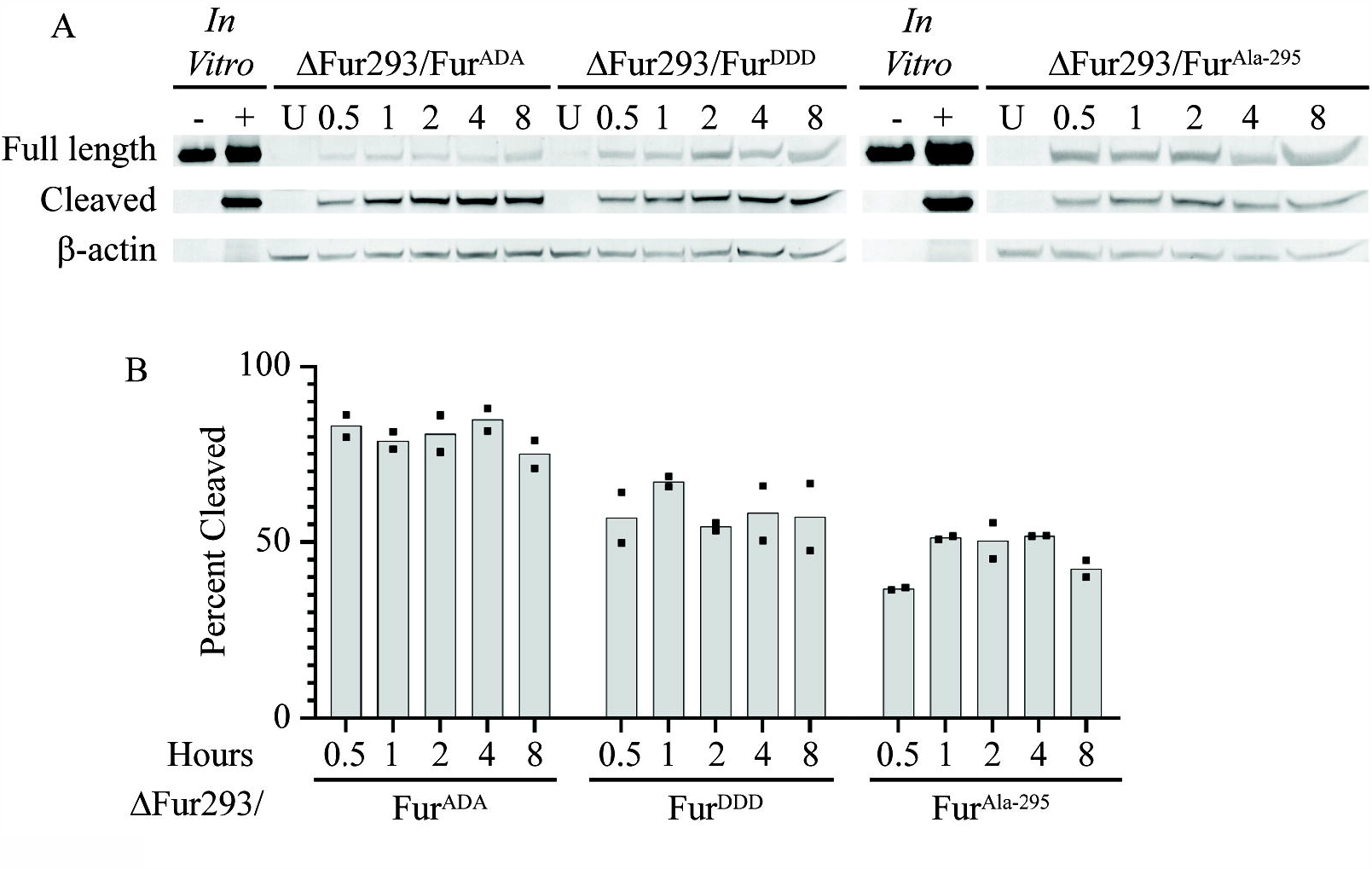
Cleavage by mutant furin. ΔFur293 cells stably expressing transgenic furin mutants (FurADA, FurDDD, and FurAla-295) were incubated for various time intervals from 0.5 to 8 hours in culture with the anti-transferrin receptor/PE24 RIT HB21-LR. Whole cell lysates were evaluated for full length and cleaved HB21-LR by western blot (panel A) and densitometry as described. Also shown are untreated (U) cell lysates for each cell line, HB21-LR in vitro with (+) and without (-) furin treatment, and the β-actin loading control. The ratio between the furin-cleaved band intensity and the total intensity of all RIT bands at each time point is plotted in panel B. The individual densitometric analysis values (points) and mean (bar) for at least two internalization and cleavage assays in each cell line are shown.

## Discussion

Here we present a furin deficient derivative of the HEK293 cell line, ΔFur293. Successful disruption of the furin gene by CRIPR/Cas9 was confirmed by western blot analysis (Figure 1) and sequencing (Supplementary Figure 1). STR profile analysis (Supplementary Table 2) confirmed the origin and identity of ΔFur293. Although variation from the reference profile was observed, the variation was consistent with existing variation in the reference sequence^36^, reports in the literature^43^, and variation among STR profiles from biological resource centers.

Cytotoxicity analysis of HEK293 FRT and ΔFur293 cells in the presence and absence of the furin inhibitor PPCI (Figure 2) showed that cells in the absence of furin or in the presence of a furin inhibitor are approximately 10-fold more resistant to HB21-LR. No cytotoxicity difference was found when comparing ΔFur293 cells to HEK293 FRT cells with PPCI. These data conclusively demonstrate that furin enhances toxin sensitivity. A small (1.5-fold) but significant difference was observed when both PPCI-treated HEK293 FRT and PPCI-treated ΔFur293 cells were evaluated for cytotoxicity. This observation suggests that cells may retain a small quantity of active furin in the presence of PPCI, and support the use of ΔFur293 over PPCI for future studies.

Insertion of the wild type furin gene into ΔFur293 cells (ΔFur293/Fur) restored and enhanced RIT sensitivity (Figure 3). This observation had not previously been observed, but is consistent with our expectation that increased furin expression can enhance RIT cytotoxicity. Our conclusion was confirmed by overexpression of transgenic furin in HEK293 FRT cells. Control experiments with transgenic chloramphenicol acetyltransferase showed no difference in RIT sensitivity from the parental line.

We next performed a RIT intracellular cleavage analysis of the HEK293 FRT, ΔFur293, and ΔFur293/Fur cell lines treated with HB21-LR (Figure 4). Compared to the parent line, ΔFur293 cells show greatly diminished cleavage and ΔFur293/Fur cells show enhanced cleavage. This is consistent with the cytotoxicity data and western blot evaluation of expression levels (Supplementary Figure 5). We further observed that ΔFur293 cells exhibited a small percentage of cleaved toxin. The presence of cleavage products in these cells may be due to other PCSKs with similar cleavage site specificities. It is also possible that cleavage sites for lysosomal proteases could result in similarly-sized protein fragments that would be indistinguishable by western blot analysis^44^. Further research will be needed to explore the influence of other proteases on RIT cleavage and cytotoxicity.

We then evaluated how complementation of furin by mutants would influence RIT cleavage and cytotoxicity. The results of our analysis of agree with previous work indicating that furin cleavage is important to RIT cytotoxicity, but is not the sole means by which furin influences the efficacy of PE-based RITs^22^. Two trafficking mutants with alterations in the cytoplasmic tail at phosphorylation signal sequences Ser^773^ and Ser^775^ (Fur^ADA^ and Fur^DDD^) and a mutant with diminished catalytic activity (Fur^Ala-295^) were also evaluated.

The S773A/S775A alanine trafficking mutant of furin (Fur^ADA^) causes it to become trapped in early endosomes^38,41^. We expected no change in cleavage because PE RITs are most likely cleaved by furin in early endosomes. Although we observed no difference between RIT cleavage in ΔFur293/Fur and ΔFur293/Fur^ADA^ cells (Figures 4 and 6), we did observe an 8-fold resistance to RITs in cells expressing Fur^ADA^ (Figure 5). These results suggest that the decrease in cytotoxicity is due to furin being trapped in early endosomes and are consistent with the hypothesis that furin has a transport role during RIT intoxication.

The S773D/S775D aspartic acid trafficking mutant of furin (Fur^DDD^) is depleted from early endosomes and preferentially retrieved to the TGN or cell surface, exhibiting a more disperse localization in the cell^38^. We expected that RIT cleavage would be decreased in cells expressing this furin variant because of the constitutive signal directing furin out of early endosomes. Consistent with our expectations, cleavage was decreased by approximately 20% when compared to both ΔFur293/Fur and ΔFur293/Fur^ADA^ cells (Figure 6). Cytotoxicity, however, was decreased nearly 3-fold compared to the wild-type furin transgene and increased 2.6-fold compared to the alanine trafficking mutant (Figure 5). This intermediate cytotoxicity phenotype indicates that the altered localization pattern of the aspartic acid trafficking mutant provides a relative improvement in cytotoxicity compared to ΔFur293/Fur^ADA^. The result is consistent with the furin acting in a transport role during RIT intoxication, as the movement of furin out of early endosomes to the TGN is enhanced.

The N295A furin oxyanion hole mutant (ΔFur293/Fur^Ala-295^) has decreased activity compared to wild type furin^40^ while having an identical trafficking pattern^42^. Our results confirmed a decrease in RIT cleavage efficiency. The ΔFur293/Fur^Ala-295^ cells are approximately 40% less efficient at cleaving the RIT when compared to ΔFur293/Fur cells (Figures 4 and 6).

This resulted in an approximately 8-fold decrease in cytotoxicity relative to ΔFur293/Fur (Figure 5). Comparing ΔFur293/Fur^Ala-295^ cells to ΔFur293/Fur^ADA^ cells demonstrated a decrease in cleavage efficiency but no difference in cytotoxicity. This indicates that interfering with either the trafficking or catalytic activity of furin have similar inhibitory effects on RIT intoxication.

We conclude that altering the trafficking pattern or blocking the activity of furin can affect RIT cytotoxicity. Our results are consistent with the hypothesis that furin acts in a transport capacity to direct PE and PE-based RITs out of endosomes and into the TGN for retrograde trafficking to the ER. Although these experiments have tested only a single cell line (HEK293) with a single RIT (HB21-LR), the results are consistent with previous evidence from other RITs on other cell lines^22^ and we believe that our results will be applicable in other cases as well. Future experiments evaluating the colocalization of furin and RITs by immunofluorescence will explore this hypothesis further.

## Acknowledgements

The authors would like to thank Ira Pastan and the Laboratory of Molecular Biology, CCR, NCI, NIH for their support and encouragement of this work. We are also grateful for helpful comments and suggestions from Margaret Regan, Matthew Bonett, Eber Guzman-Cruz, and Dayshia Kerney.

## Funding and additional information

This work was supported by funding from Towson University, including a Faculty Development & Research Committee award for JEW, awards from the Office of Undergraduate Research and Creative Inquiry and the Jess and Mildred Fisher College of Science and Mathematics for JDS and YZ, and awards from the Graduate Student Association for BDG and YZ.

## Footnotes

## Abbreviations used

RIT: recombinant immunotoxin,
PE: *Pseudomonas* exotoxin
A, PCSK: proprotein convertase subtilisin/kexin,
FRT: Flp recombinase recognition target site, and sg
RNA: single guide RNA.

