## Supplementary Figures and Tables for "Intracellular trafficking of furin enhances cellular intoxication by recombinant immunotoxins based on *Pseudomonas* exotoxin A"

*Title*

*Contents*

Figure S1. ∆Fur293 cells contain a deletion in exon 2 of the furin gene

Figure S2. PPCI diminishes the toxicity of HB21

Figure S3. Representative HB21 cytotoxicity assays

Figure S4. Cytotoxicity evaluation of PPCI-treated LoVo cells

Figure S5. Furin expression levels

Table S1. Oligonucleotides

Table S2. STR profiles

Table S3. Statistical analysis of normalized EC_50_ differences between furin transgenic cell lines

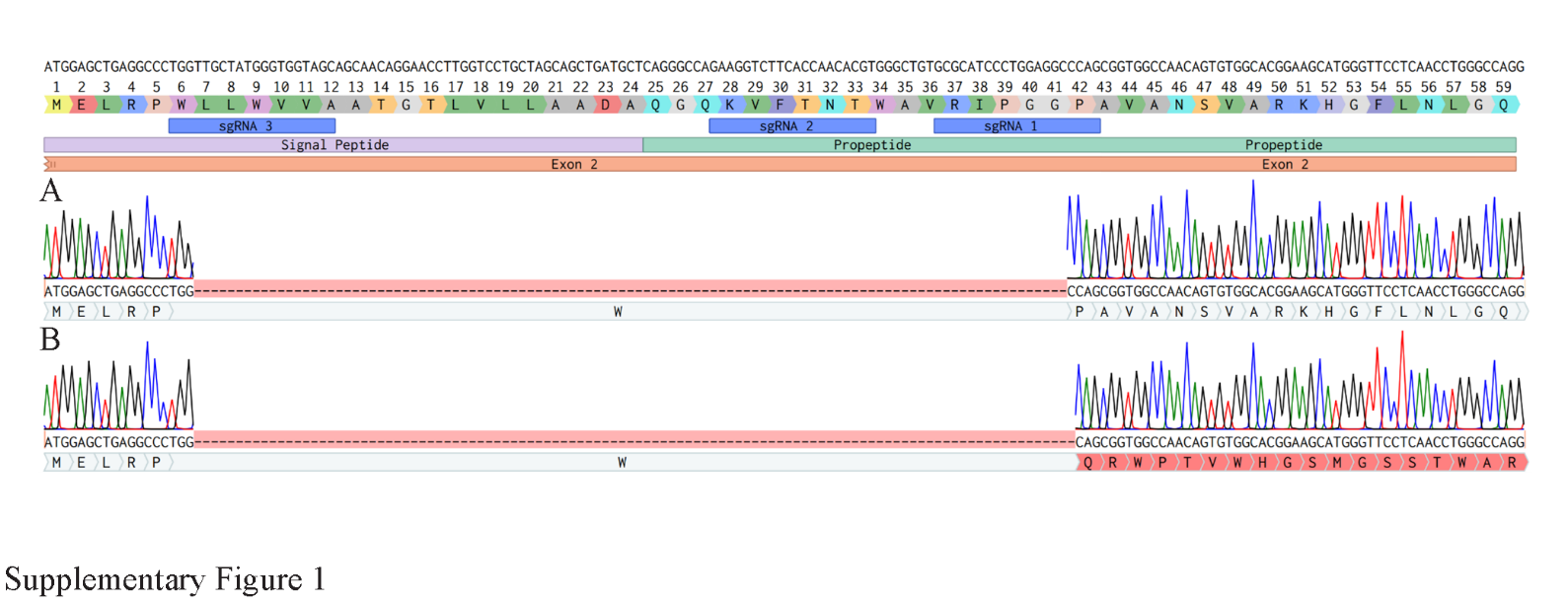
*Figure S1. ∆Fur293 cells contain a deletion in exon 2 of the furin gene.* Alignment of exon 2 from human furin with sequencing results from ∆Fur293. Panel A shows a representative chromatogram of the 105 bp deletion. Panel B shows a representative chromatogram of the 106 bp deletion. Images were generated using Benchling^1^.

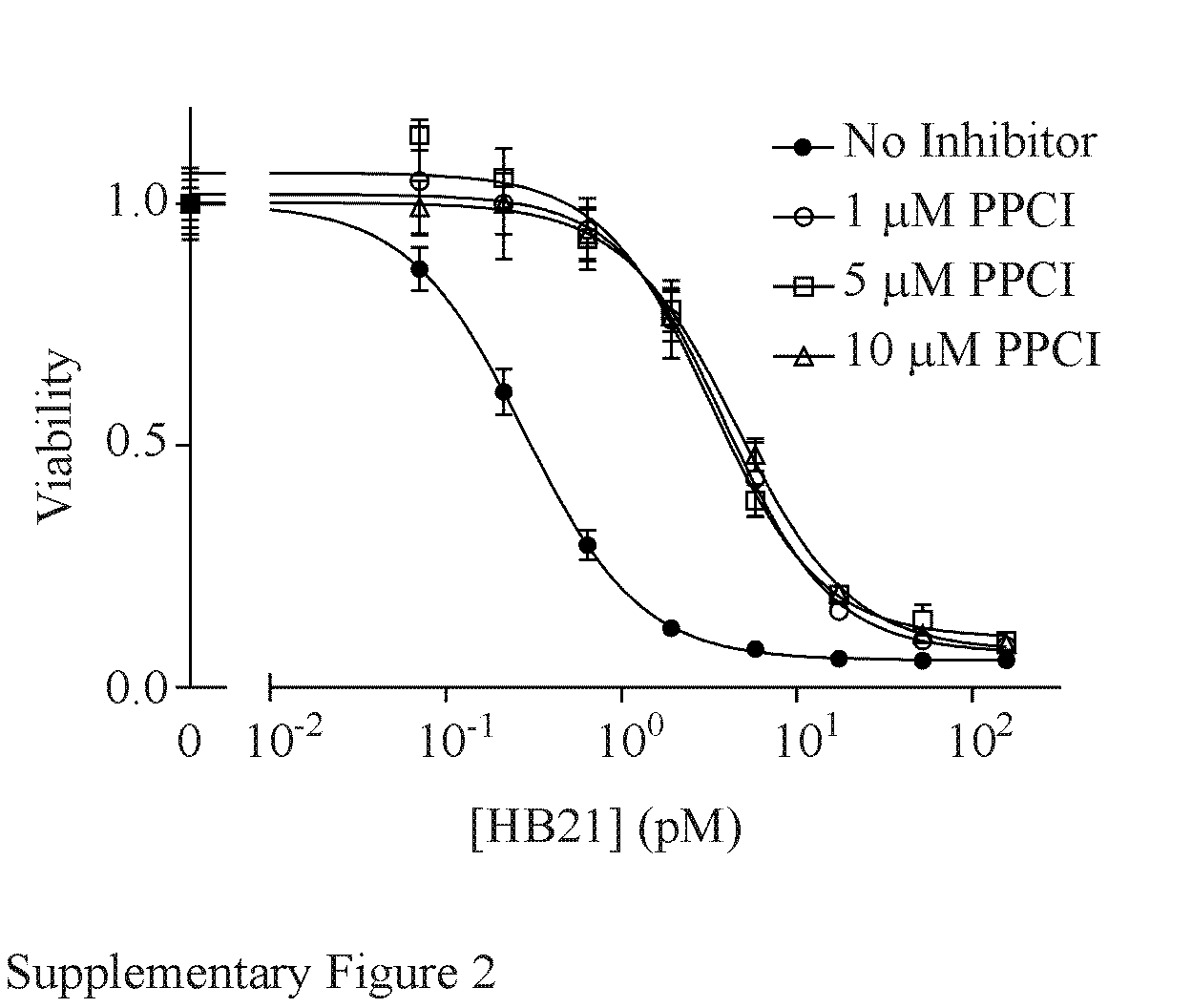

*Figure S2. PPCI diminishes the toxicity of HB21.* The cytotoxicity of HB21 was evaluated against HEK293 FRT cells in the presence of increasing concentrations of proprotein convertase inhibitor (PPCI). Error bars indicate the standard error of the mean for a minimum of 5 replicates at each toxin concentration. (No Inhibitor EC_50_ = 0.2806 pM; 1 μM PPCI EC_50_ = 3.923 pM; 5 μM PPCI EC_50_ = 3.263 pM; 10 μM PPCI EC_50_ = 4.557 pM).

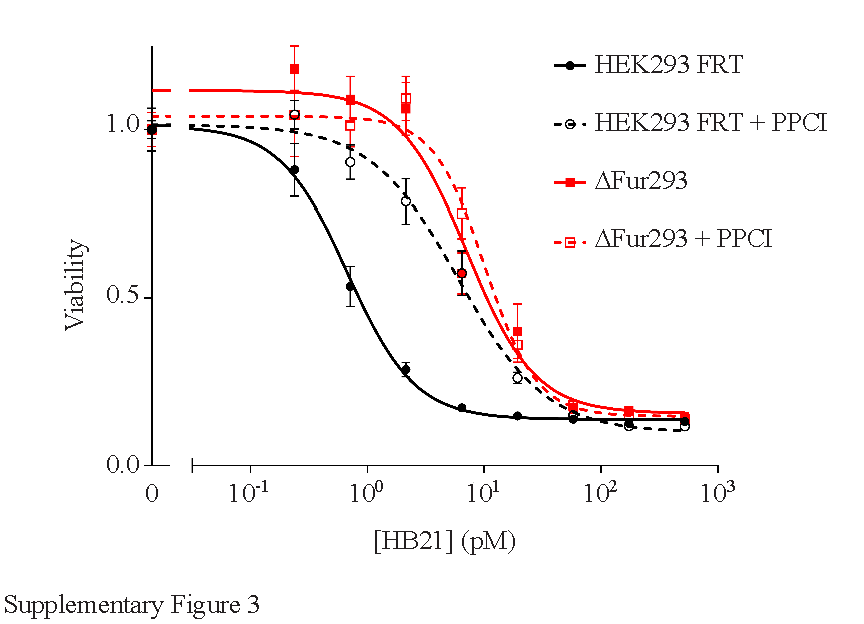

*Figure S3. Representative HB21 cytotoxicity assays.* The cytotoxicity of HB21 was evaluated against HEK293 FRT and ∆Fur293 cells in the absence and presence of 1 μM PPCI. Error bars denote standard error values for a minimum of 5 replicates at each toxin concentration.

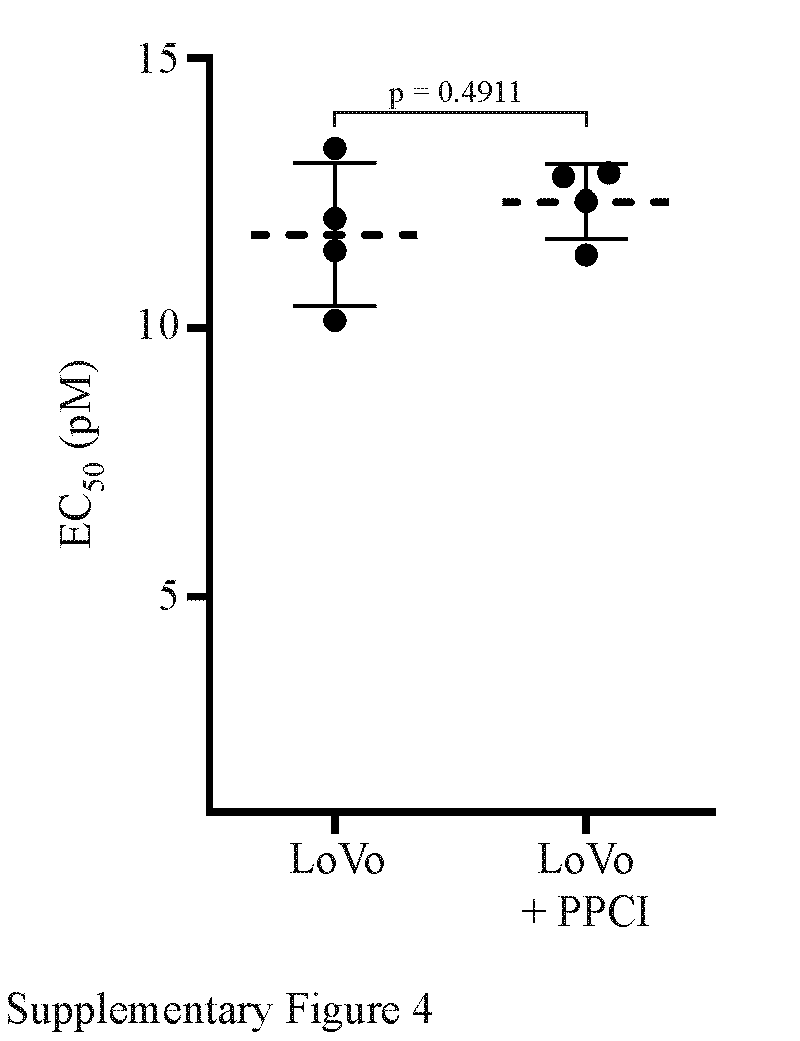

*Figure S4. Cytotoxicity evaluation of PPCI-treated LoVo cells.* Furin-deficient LoVo cells were treated with HB21 in the presence and absence of 1 μM PPCI. EC_50_ (pM) values from four separate paired assays are plotted. Dotted lines denote the average value for each condition and error bars indicate the standard deviation. No significant difference was detected using a T-test performed as described.

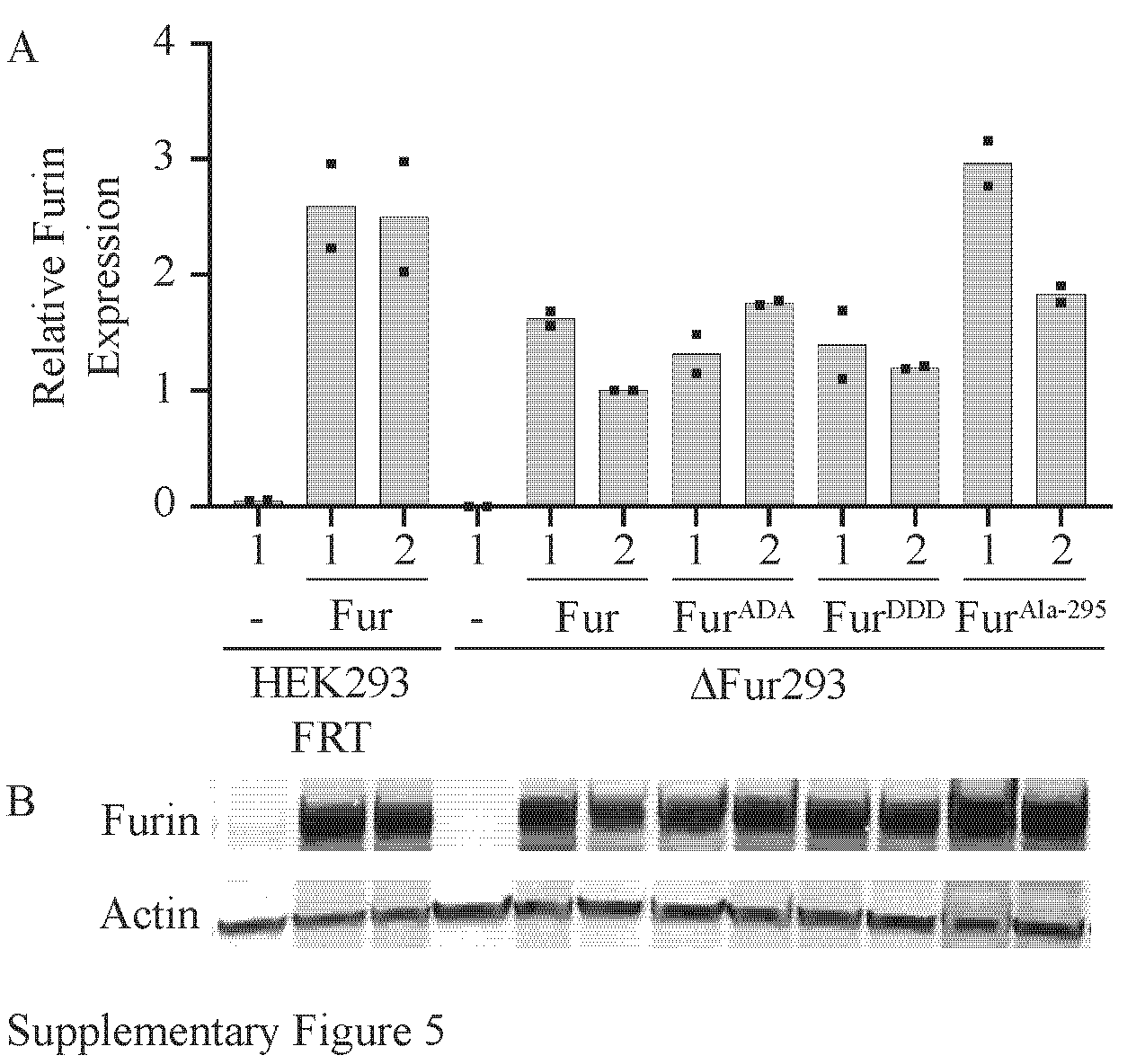

*Figure S5. Furin expression levels.* Cell lysates from HEK293 FRT and ∆Fur293 cell lines with and without transgenic furin (Fur, Fur^ADA^, Fur^DDD^, or Fur^Ala-295^) were evaluated for furin expression by western blot. Two separate clones from each transgenic line were evaluated, and each line was evaluated twice. One representative western blot (panel B) is shown. Band intensity from each western blot was calculated and normalized. Results are plotted as the average of two experiments (panel A). Normalized expression values were used to correct for differences in response to HB21 cytotoxicity.

*Table S1. Oligonucleotides*

| **Oligonucleotide** | **Sequence** |
| --- | --- |
| Furin CRISPR sgRNA1 | 5’GCG CAT CCC TGG AGG CCC AG |
| Furin CRISPR sgRNA2 | 5’GAA GGT CTT CAC CAA CAC GT |
| Furin CRISPR sgRNA3 | 5’GCT ACC ACC CAT AGC AAC CA |
| pCMV6 Insert Start For | 5’TCT AGT AAG CTT GCC GCC GCG ATC GCC ATG |
| pCMV6 Insert End Rev | 5’GCC GGC TTA AAC CTT ATC GTC GTC |
| T7 Forward | 5’TAA TAC GAC TCA CTA TAG GG |
| CMV Forward | 5’CGC AAA TGG GCG GTA GGC GTG |
| hEF2 Seq Primer Reverse | 5’GAT GGC TGG CAA CTA GAA GGC AC |
| BGH Reverse | 5’TAG AAG GCA CAG TCG AGG |
| hFUR SDS>ADA F [BM] | 5’ACG CAG AAG AGG ACG AGG GCC G |
| hFUR SDS>ADA R [BM] | 5’CAG CCG GCC ACT CCT CCT GCC A |
| hFUR SDS>DDD F [BDG] | 5’ACG ATG AAG AGG ACG AGG GCC G |
| hFurin CDNA Truncated R | 5’TGG CAT CGT ACG CGT GGC CTG ACT GGA CGT GAG |
| hFUR Exon 1 3 PCR For | 5’CAT CTG TCC ATC TAT GTG GCT GGC |
| hFUR Exon 1 3 PCR Rev | 5’GCA GGA GGA AAG GAA ACG AAT CAG AG |
| hFur Exon 1 Seq | 5’GCA GGG CAG CTT TAG TGC GTA G |
| pUC57 Furin Ex 2 OvEx C+A | 5’CCG CAT CAC CAT CAT CAC CAC TAA GAA TTC ATC GCA GGG CAG CTT TAG TGC GTA G |
| pUC57 Furin Ex 2 OvEx D+B | 5’GCG GAT AAC AAT TTC ACA CAG GAA ACA GCT ATG ACC TCA CTC TCA CTG GGA TAT GCT GC |
| hFurin cDNA Catalytic Domain Sequencing F | 5’GTG GCA AAG CGA CGG ACT AAA C |
| hFurin cDNA P Domain R primer | 5’GAT GTC GAT GAT GCA CTT CCG C |
| hFurin cDNA P Domain F | 5’GCG GAA GTG CAT CAT CGA CAT C |
| hFurin cDNA Catalytic Domain R | 5’GTT TAG TCC GTC GCT TTG CCA C |
| M13 R | 5’CAG GAA ACA GCT ATG AC |

*Table S2. STR profiles*

| **Locus** | **HEK293^†^** | **HEK293 FRT** | **HEK293 ΔFUR** |
| --- | --- | --- | --- |
| Amelogenin | X^2–11^ | X | X |
| CSF1PO | 12*^10^  11,12^4–9,11^ | 12 | 12 |
| D1S1656 | *Not reported* | 15,17.3 | 15,17.3 |
| D2S441 | *Not reported* | 11,15 | 11,15 |
| D2S1338 | 19^8^ | 19 | 19 |
| D3S1358 | 15*^2^  15,17^8,9^ | 15,17 | 15,17 |
| D5S818 | 8*^10^  8,9^2,4–9,11^ | 8 | 8 |
| D7S820 | 11,12^2,4–11^ | 11,12 | 11,12 |
| D8S1179 | 12,14^2,3,8^ | 12 | 12,14 |
| D10S1248 | *Not reported* | 14 | 14 |
| D12S391 | *Not reported* | 19,21 | 19,21 |
| D13S317 | 12^4^  12,14^2,5–11^ | 12,14 | 12,14 |
| D16S539 | 9  9,13^4–8,10,11^ | 9,13 | 9,13 |
| D18S51 | 17^2,3^  17,18^3^  18^8^ | 17,18 | 17,18 |
| D19S433 | 15,18  18^8^ | 15,18 | 18 |
| D21S11 | 28,30.2^2,3,8^ | 28,30.2 | 28,30.2 |
| D22S1045 | *Not reported* | 16 | 16 |
| FGA | 23^2,3,8,9^ | 23 | 23 |
| Penta D | 9,10 | *Not evaluated* | *Not evaluated* |
| Penta E | 7,15 | *Not evaluated* | *Not evaluated* |
| SE33 | *Not reported* | 17 | 17 |
| TH01 | 7,9.3^3–11^ | 7,9.3 | 7,9.3 |
| TPOX | 11^4–11^ | 11 | 11 |
| vWA | 16*^3^  16,19^2–11^ | 16,19 | 16,19 |

**^†^**STR data as reported for HEK293 (RRID:CVCL_0045) by the *Cellosaurus* database^12^. Note that multiple variants were reported for several loci. Additional variants not listed in the *Cellosaurus* database (indicated by *) were also noted during a search of literature and biological resource centers. Not all loci could be corroborated, but references are provided where possible.

*Table S3. Statistical analysis of normalized EC_50_ differences between furin transgenic cell lines*

|  | ΔFur293 | Fur  #1 | Fur  #2 | Fur^ADA^  #1 | Fur^ADA^  #2 | Fur^DDD^  #1 | Fur^DDD^  #2 | Fur^Ala-295^  #1 | Fur^Ala-295^  #2 |
| --- | --- | --- | --- | --- | --- | --- | --- | --- | --- |
| ΔFur293 |  |  |  |  |  |  |  |  |  |
| Fur #1 | 0.0150 |  |  |  |  |  |  |  |  |
| Fur #2 | 0.0152 | 0.1423 |  |  |  |  |  |  |  |
| Fur^ADA^ #1 | 0.0134 | 0.0082 | 0.0052 |  |  |  |  |  |  |
| Fur^ADA^ #2 | 0.0126 | 0.0322 | 0.0240 | 0.8619 |  |  |  |  |  |
| Fur^DDD^ #1 | 0.0284 | 0.0129 | 0.0137 | 0.0035 | 0.0138 |  |  |  |  |
| Fur^DDD^ #2 | 0.0140 | 0.0064 | 0.0010 | 0.0029 | 0.0192 | 0.3624 |  |  |  |
| Fur^Ala-295^ #1 | 0.0395 | 0.0383 | 0.0291 | 0.2877 | 0.2994 | 0.0201 | 0.0232 |  |  |
| Fur^Ala-295^ #2 | 0.0129 | 0.0011 | 0.0004 | 0.5947 | 0.6885 | 0.0003 | 0.0003 | 0.9999 |  |
